# Numerical investigation of the dynamics of a rigid spherical particle in a vortical cross-slot flow at moderate inertia

**DOI:** 10.1101/2022.12.19.520995

**Authors:** Konstantinos Kechagidis, Benjamin Owen, Lionel Guillou, Henry Tse, Dino Di Carlo, Timm Krüger

**Affiliations:** School of Engineering, Institute for Multiscale Thermofluids, University of Edinburgh, Edinburgh, United Kingdom; Cytovale, Inc., San Fransisco, California, United States of America; Department of Bioengineering, University of California, Los Angeles, California, United States of America

**Keywords:** Vortex flow, cross-slot flow, inertial microfluidics, particle dynamics, lattice-Boltzmann method

## Abstract

The study of flow and particle dynamics in microfluidic cross-slot channels is of high relevance for lab-on-a-chip applications. In this work we investigate the dynamics of a rigid spherical particle in a cross-slot junction for a channel height-to-width ratio of 0.6 and at a Reynolds number of 120 for which a steady vortex exists in the junction area. Using an in-house immersed-boundary-lattice-Boltzmann code, we analyse the effect of the entry position of the particle in the junction and the particle size on the dynamics and trajectory shape of the particle. We find that the dynamics of the particle depends strongly on its lateral entry position in the junction and weakly on its vertical entry position; particles that enter close to the centre show trajectory oscillations. Larger particles have longer residence times in the junction and tend to oscillate less due to their confinement. Our work contributes to the understanding of the particle dynamics in intersecting flows and enables the design of optimised geometries for cytometry and particle manipulation.

## 1 Introduction

The deformability of blood cells is a valuable label-free biomarker to indicate changes in the cell state [1]. Changes in the nucleus or cytoskeleton of blood cells are associated with various diseases, including malaria [2, 3], sickle cell disease [4], cancer [5] and sepsis [6]. For example, oral cancer cells have been found to be more deformable than healthy oral epithelial cells [7], thus facilitating metastasis [8]. Leukocytes have been shown to become stiffer or slower passing through pores upon stimulation with chemokines associated with infection, such as sepsis [9]. Conventional methods for single cell analysis, for instance, atomic force microscopy [10], optical stretching [11] and micropipette aspiration [12] are useful for fundamental research in mechanobiology, but their low throughput limits the potential for clinical applications.

Fast and accurate analysis of blood samples is important for identifying diseased cells, thus accelerating clinical decision making [13]. Inertial microfluidics (IMF) has been widely used for cell manipulation, separation and characterisation in biomedical applications [14]. IMF operates in a flow regime where inertial forces influence the fluid and particle dynamics [15]. In the 1960s, Segré and Silberberg [16, 17] observed radial migration of rigid spherical particles towards an annulus at around 60%of the radius of a tube when the Reynolds number is of the order 10–100. Since then, several studies investigated the nature of the inertial lift forces in straight channels [18–20]. In the 2000s, it was recognised that the Segré-Silberberg effect could be exploited in a microfluidic setting [21] and in more complex channel geometries, such as curved [22, 23], serpentine [24, 25] and spiral channels [26–28].

Since its inception, IMF has played an important role in image-based deformability cytometry (DC). DC uses different channel geometries, such as linear channels [29], constrictions [30] and cross-slot channels [31]. While traditional cytometry techniques, such as atomic force microscopy, micropipette aspiration and optical stretchers, are limited to low cell processing rates (10–100 cells/hr) [29], DC yields significantly higher throughput (around 10–1000 cells/sec) [32], making it more suitable for diagnostic applications where large quantities of cells should be measured to have reliable measurements.

Cross-slot channels (Fig. 1) are particularly appealing for DC since the flow field in their junction features both high stresses, therefore deforming cells, and a relatively low flow speed, thus facilitating optical imaging. The fluid flow in cross-slot junctions is complex and has been extensively investigated. For specific ranges of channel aspect ratio and Reynolds number, a spiral vortex can appear in the junction [33] which has been explained by a symmetry-breaking bifurcation above a critical Reynolds number for a given channel aspect ratio [34]. Furthermore, Burshtein et al. [35] determined the characteristics of a steady-state vortex in cross-slot flows at higher Reynolds numbers, such as the vortex core structure and time-periodic fluctuations around the stagnation point in the junction, by tuning the aspect ratio of the device. The design of the junction has been optimised to achieve more homogeneous strain rates [36, 37] while Zhang *et al.* [38] identified different flow regimes in the cross slot by changing the flow conditions. Recently, a cross-slot device has aided the early detection of sepsis by using the deformability of leukocytes as a biomarker [39].

**Figure 1:**
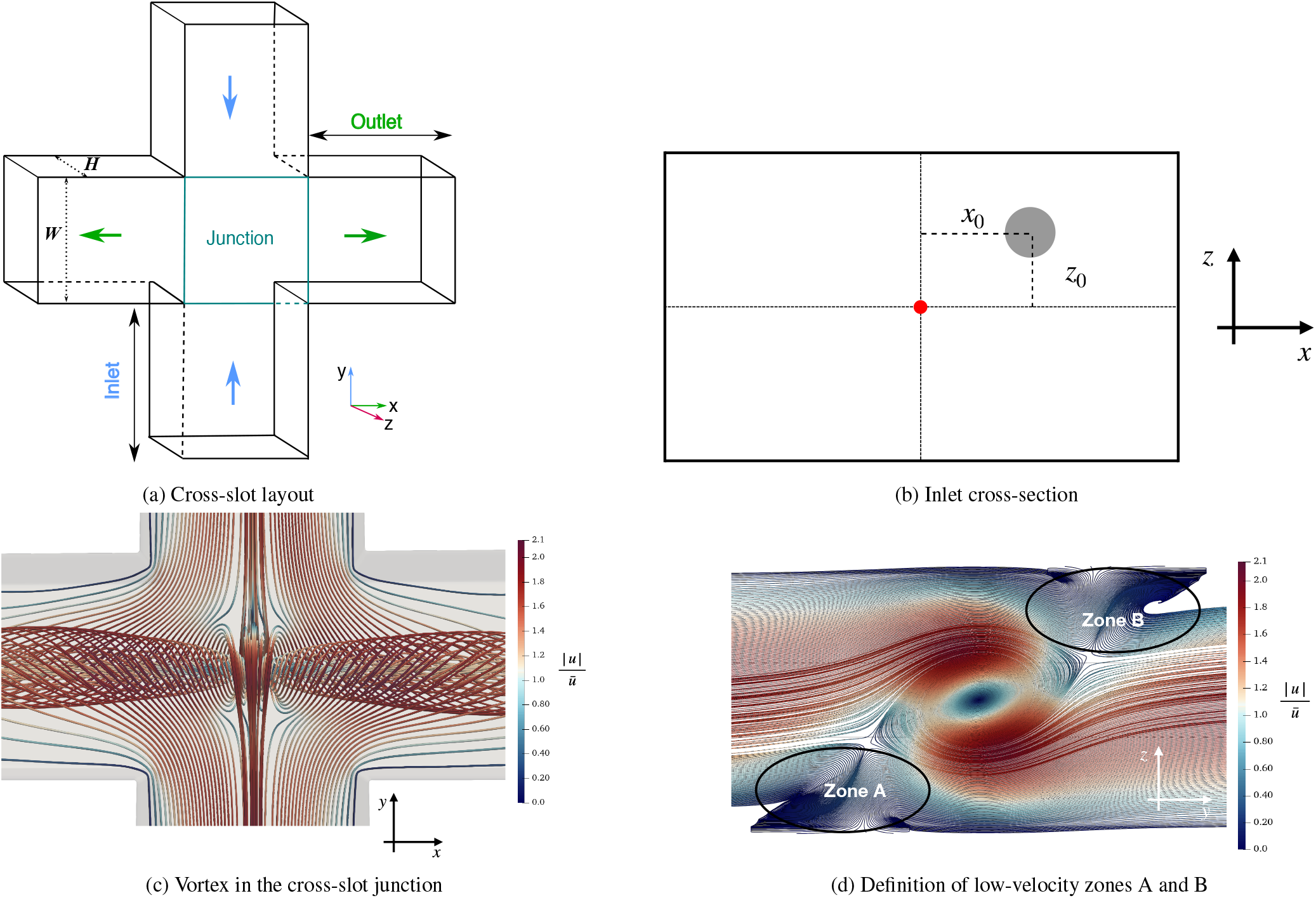
(a) 3D illustration of the cross-slot geometry with two inlets and two outlets. (b) Definition of the initial position (*x*_0_, *z*_0_) of the particle’s centre of mass on the inlet cross-section. The origin (red dot) is located on the inlet’s centre-line, and the particle initially moves along the *y*-axis (pointing into the plane of view). (c) Top view of streamlines showing the spiral vortex at *Re* = 120 and channel aspect ratio *α* = 0.60, as defined in Sec. 2.1. The colour bar indicates the velocity magnitude (normalised by the average inlet velocity). (d) Definition of low-velocity zones A and B in the junction which are important for the particle dynamics. View along the outlet axis.

Despite earlier works focused on the fluid dynamics and particle trapping in cross-slot junctions, the interaction of a particle and the vortex at moderate inertia has not been thoroughly investigated. It is currently unclear how flow field characteristics and particle properties correlate with observable metrics, such as the shape of the particle trajectory and the residence time in the junction. In this paper, we use immersed-boundary-lattice-Boltzmann simulations to explore the dynamics of a spherical, rigid particle in an incompressible Newtonian liquid in the steady-vortex regime (Sec. 2). We pay particular attention to the characterisation of the geometry, observables and simulation initialisation (Sec. 3). We investigate the effect of initial particle position and size on the trajectory and residence time of the particle in the cross-slot junction at a Reynolds number of 120 and a channel aspect ratio of 0.6 (Sec. 4). Our results show that particle trajectory and residence time strongly depend on the initial lateral position. Under the investigated flow conditions and channel geometry, the vortex affects larger particles less compared to smaller ones. In Sec. 5, we provide a concluding discussion. We hope that our results stimulate further work, for example the investigation of deformable particles in the cross-slot junction.

## 2 Physical model and numerical methods

We briefly outline the physical model (Sec. 2.1) and the numerical methods (Sec. 2.2) used in this work.

### 2.1 Physical model, parameters and non-dimensional groups

We consider a single rigid, spherical particle immersed in an incompressible, isothermal, viscous Newtonian liquid. The incompressible Navier-Stokes equations and Newton’s laws of motion determine the dynamics of the liquid and the particle, respectively. The no-slip condition is assumed at all solid-liquid boundaries.

The relevant parameters characterising the fluid flow are the liquid density *ρ*_f_, the kinematic viscosity *ν*, and the mean velocity *ū* in the inlet of the cross-slot geometry. The particle is fully characterised by its radius *a* and density *ρ*_p_. As will be described in more detail in Sec. 3, the geometry is fully determined by the width *W* and height *H* of the inlet and outlet channels of the cross-slot geometry.

Following [34], we define the channel Reynolds number as

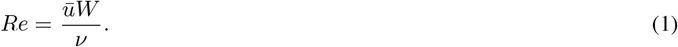

The non-dimensional confinement of the particle in the cross-slot geometry is

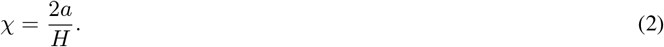

The aspect ratio of the cross-section of the inlet and outlet channels is given by

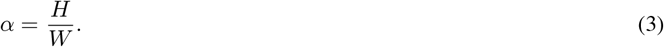

Note that we chose *α* < 1 in our study, justifying the use of *H* in the definition of the confinement, Eq. (2). The hydraulic diameter

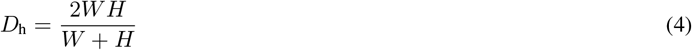

is important for the characterisation of the inflow behaviour of the particle. We further define the advection time scale

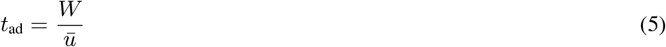

to non-dimensionalise time as 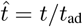.

### 2.2 Numerical methods

We employ the lattice-Boltzmann (LB) method with the D3Q19 lattice [40], the BGK collision operator [41] and the Guo forcing scheme [42] to solve the Navier-Stokes equations. The relaxation time *τ* of the BGK collision operator is linked to the kinematic viscosity according to

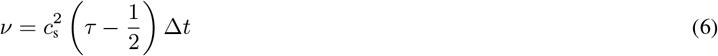

where 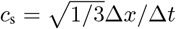 is the lattice speed of sound, Δ*x* is the lattice spacing and Δ*t* is the time step.

The multi-direct-forcing immersed-boundary method with a three-point stencil is used to satisfy the no-slip boundary condition on the surface of the moving rigid particle [43, 44]. The half-way bounce-back scheme [45] is employed to satisfy the no-slip condition on the stationary walls of the cross-slot geometry. Additionally, due to non-negligible particle inertia, a coefficient of restitution is included to model perfectly elastic collisions to avoid any overlap between the particle and the wall.

We impose a fully developed rectangular Poiseuille flow with the desired mean velocity *ū* at both inlets [46]. The boundary condition at both outlets is based on an extrapolation approach [47] modified in such a way that the average pressure on the outlet plane is constant in time. Therefore, the inlet velocity and the outlet pressure are imposed, while the outlets permit the swirling flow caused by the vortex at the centre of the cross-slot geometry to leave the domain without undesired reflections at the outlet planes.

To ensure that our simulation code accurately captures the fluid-particle interaction, we compared the lateral equilibrium positions in a square cross-section with published results [48] (App. A.1).

## 3 Cross-slot geometry and simulation set-up

The geometry consists of a cross-slot channel as shown in Fig. 1a. The origin of the Cartesian coordinate system is at the centre of the cross-slot geometry. The flow in the inlets is along the *y*-axis (inlet axis), the flow in the outlets is along the *x*-axis (outlet axis), and the *z*-axis denotes the height direction (height axis). Depending on the combination of Reynolds number, height-to-width aspect ratio and ratio of the outlet flow rates, different flow regimes inside the cross-slot junction can be observed, for example a vortex which might be steady or unsteady [35,38,49]. In this work, we are interested in the steady-vortex regime since the steady vortex leads to reproducible and non-trivial particle behaviour, which is relevant for flow cytometry applications. Therefore, we set the Reynolds number to *Re* = 120, the channel aspect ratio to *α* = 0.6, and the ratio of both outlet flow rates to 1, which leads to the formation of a steady vortex [34].

Since both inlet flow rates and both outlet flow rates are identical, the resulting flow field, including the vortex, is symmetric with respect to the inlet-height plane (*yz*-plane) and under a 180°-rotation about the outlet axis (*x*-axis) in the absence of the particle. Therefore, it is sufficient to enter the particle only on one side of the inlet-height plane, and all initial particle positions are *x*_0_ > 0. Furthermore, we only inject particles at one inlet (*y*_0_ < 0). Due to the presence of the vortex spiralling about the outlet axis, we need to explore the entire height range of initial particle positions. This procedure allows us to scan all physically unique initial particle positions without running multiple simulations that are physically equivalent.

The height *H* is constant within the entire domain, and both inlets and both outlets have the same width *W*. We use 36 grid points for *H* and 60 grid points for *W*, giving 45 grid points for the hydraulic diameter *D_h_*. The length of the inlets and outlets is *L*_in_ = *L*_out_ = 4*D*_h_, which we found to be sufficient to give results independent of the inlet and outlet lengths. The relaxation time is chosen as *τ* = 0.55Δ*t* in all simulations, resulting in a viscosity of *ν* = 1/60 Δ*x*^2^/Δ*t*. The peak and average values of the inlet velocity profile are *u*_max_ = 0.068 Δ*x*/Δ*t* and *ū* = 0.033 Δ*x*/Δ*t*, respectively. We consider particle sizes resulting in confinement values *χ* in the range [0.35, 0.65]. The particle is neutrally buoyant in all cases, *ρ*_p_/*ρ*_f_ = 1.

To reduce the required time for the vortex to form, we introduce a force field that accelerates the fluid in the junction’s upper and lower halves in opposite directions along the inlet axis for a finite amount of time. Once the force has been switched off, the vortex flow is allowed to converge to a steady state with a convergence criterion of 10^-6^ for the relative difference of the flow field between consecutive time steps. It is reported that the right or left-handed orientation of the spiral vortex occurs with the same probability [34]. For consistency, we initialise the simulations in such a way that the vortex is always left-handed, as shown in Fig. 1d. Eventually, the particle is injected at the desired initial position next to the inlet at *y*_0_ < 0.

We always initialise particles with a finite value of *x*_0_ since a particle initially on the mid-plane between the outlets would be caught in the vortex indefinitely in the absence of distortions. Since numerical noise in the simulation would eventually let the particle escape in one of the outlets, the residence time would be determined by numerical noise and not deterministic physics. In real-world experiment, channel and particle imperfections always lead to finite residence times in the geometry considered here.

Due to inertial forces, particles initialised near the inlet would migrate away from their prescribed lateral coordinates (*x*_0_, *z*_0_) while moving along the inlet, therefore losing control over the location where the particle enters the actual junction. To avoid this premature particle migration in the inlet, we lock the x- and z-components of the particle position while allowing the particle to rotate and move along the inlet axis freely. The ‘guidance’ of the particle is switched off when the particle reaches a point located *D*_h_ upstream of the junction; the particle is allowed to move freely afterwards. Our simulations show that, at an upstream distance of about one hydraulic diameter from the junction, the velocity profile in the inlet starts to be affected by the junction. Therefore, the initial cross-sectional position (*x*_0_, *z*_0_) defines at which point the particle enters the region of influence of the junction.

In the interest of using a concise notation, we define

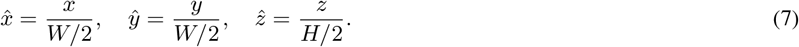

Therefore, the accessible range of initial positions for a point particle in the inlet channel is 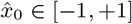 and 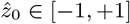.

## 4 Results and discussion

We present and discuss our simulation results in two main sections. Sec. 4.1 covers the effect of initial particle position 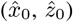 on the trajectory and residence time of the particle. Sec. 4.2 focuses on the effect of particle size, including a neutrally buoyant point particle.

### 4.1 Effect of initial particle position

To study the effect of the initial particle position, we investigated a range of initial positions 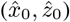 as summarised in Tab. 1. A first simulation study revealed that the richest particle dynamics can be found for small values of 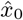. Hence, we ran more subsequent simulations for smaller values of 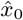. For the values of 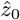, we chose a starting position on the mid-plane 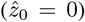 and the equilibrium positions of differently sized particles (*χ* between 0.35 and 0.75) in a straight duct with the same aspect ratio (*α* = 0.60) as the inlet of the cross-slot geometry at *Re* = 120, see App. A.2. The same confinement values are used in Sec. 4.2 to investigate the effect of particle size. The particle is neutrally buoyant and has a fixed confinement of *χ* = 0.4. The channel aspect ratio is *ρ* = 0.6, and the Reynolds number is 120.

**Table 1:** Non-dimensional initial particle positions 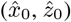 used in Sec. 4.1. See Fig. 1b and Eq. (7) for the definition of initial positions.

|  |  |
| --- | --- |
| $\hat{x}_0$ | 0.0033, 0.01, 0.05, 0.075, 0.15, 0.25, 0.50 |
| $\hat{z}_0$ | 0, $\pm 0.22$, $\pm 0.35$, $\pm 0.43$, $\pm 0.49$ |

#### 4.1.1 Particle trajectories

Fig. 2a and 2b show some representative particle trajectories inside the cross-slot junction for two different values of 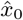 and a range of values of 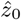. The trajectories of rigid particles found here are qualitatively similar to those presented by Gosset *et al.* [31] in their supporting videos and Zhang et *al.* [50].

**Figure 2:**
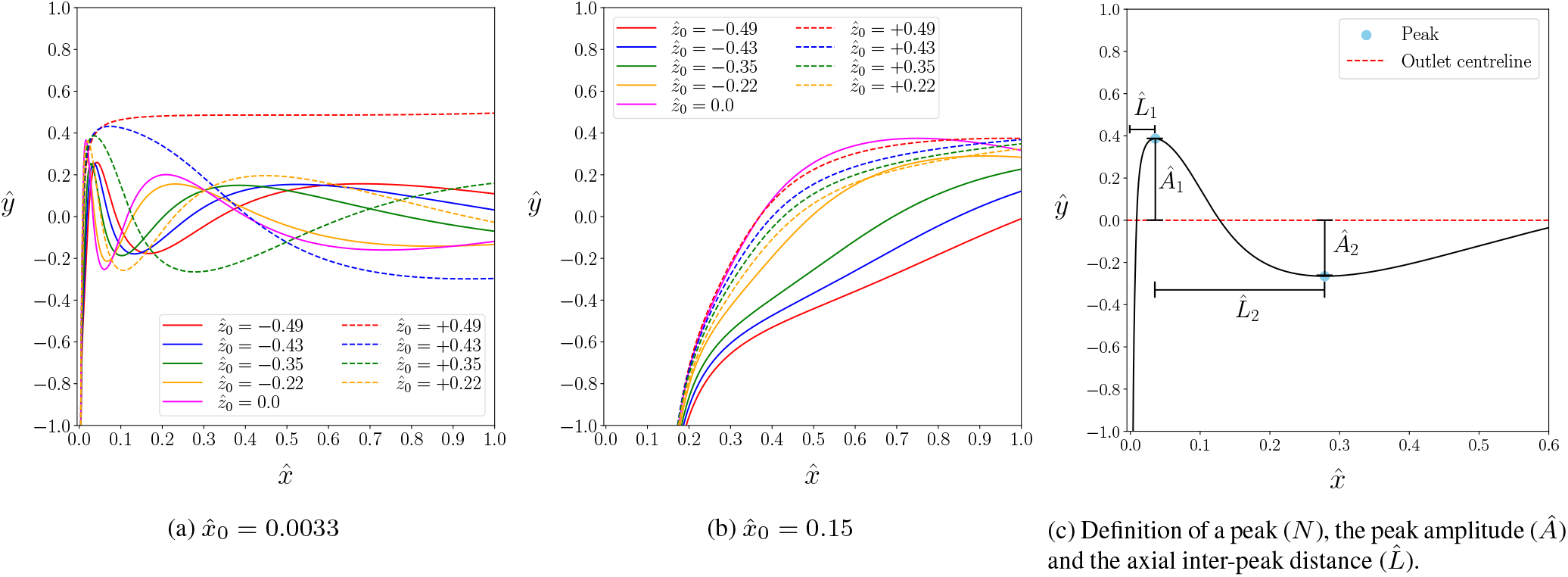
Particle trajectories projected onto the inlet-outlet plane (top view). The channel aspect ratio is *α* = 0.6, the Reynolds number is *Re* = 120, and the particle confinement is *χ* = 0.4. Different curves show different values of 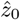 for (a) 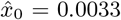 and (b) 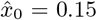. Solid and dashed lines refer to negative and positive values of 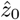, respectively (c) Illustration of a typical particle trajectory. Peaks (blue dots) are extrema of the curve, *Â_i_* denotes the *i*-th amplitude (normalised by *W*/2) of a peak with respect to the centre-line of the outlet (*x*-axis), and 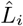 is the axial distance (normalised by *W*/2) between the (*i* – 1)-st and *i*-th peaks (with 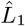 playing a special role, denoting the distance between the first peak and the centre-line of the inlet). Note that the aspect ratio of the plots has been changed to improve readability.

It is inconvenient to show the particle trajectories for all the simulations. Instead, we extract key characteristics from the particle trajectories that are also experimentally relevant. Fig. 2c illustrates the chosen ‘macroscopic’ metrics of the inlet-outlet projection (*xy*-projection, ‘top view’) of the particle trajectories: the number *N* of peaks observed in the central area of the cross-slot junction, the amplitude *A* of each peak, and the lateral distance *L* between consecutive peaks. A peak is defined as a minimum or maximum of the inlet-outlet projection of the particle trajectory. The residence time *T*_res_ is the length of time the centre of mass of the particle spends in the central junction area, 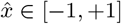 and 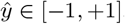.

Particles initialised very close to the mid-plane of the inlet, 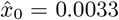 in Fig. 2a, give rise to damped oscillations in the trajectories. For moderate distances from the mid-plane, 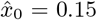 in Fig. 2b, the oscillations are either weak or completely absent. Fig. 2 also shows that 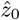 has a stronger effect on the shape of the trajectories when 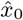 is small. We found that particles only show appreciable oscillations when the initial position 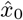 is smaller than 0.1. We will discuss these findings in the remainder of this section by providing further ‘microscopic’ information: instantaneous particle velocities (translational and angular), and forces (lift and drag) which we use to rationalise the behaviour of the macroscopic observables.

In the following, we will focus on an analysis of the macroscopic metrics defined in Fig. 2c (number of peaks, peak amplitude, inter-peak distance) and the residence time to characterise the particle trajectories and their dependence on initial positions.

#### 4.1.2 Number of peaks of particle trajectories

Fig. 3a shows the number of peaks, *N*, for all investigated initial positions 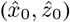. Only for the narrow range 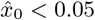, some trajectories have two or more peaks. Increasing 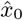 from 0.0033 to 0.01 reduces *N* by one for all the investigated values of 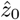. Upon a further increase of 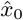, the number of peaks reduces to one or zero, depending on 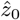. No peaks are observed for any trajectory with 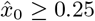.

**Figure 3:**
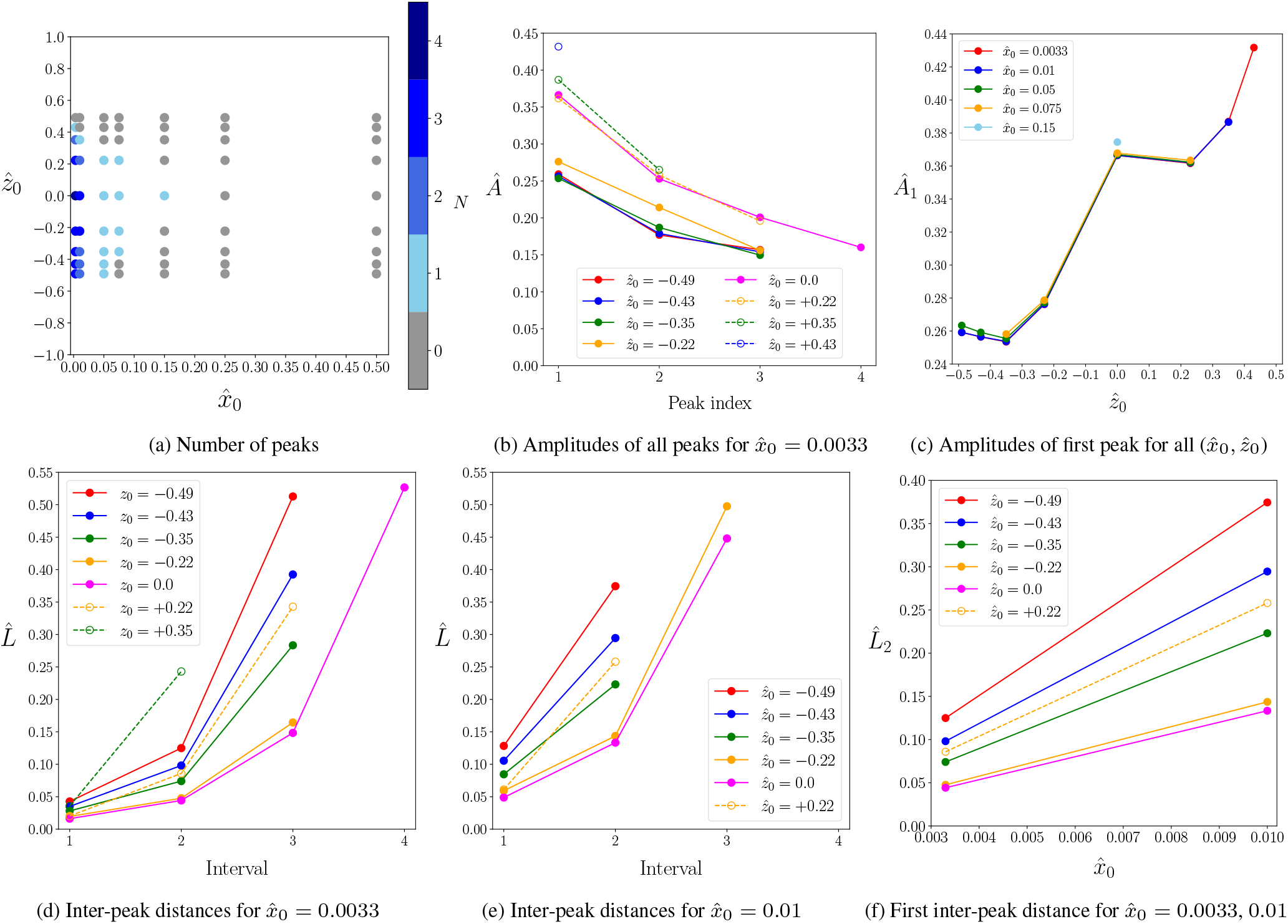
(a) Number of peaks, *N* (as defined in Fig. 2c), of the trajectories for various initial positions 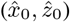. Note that both axes have different scales to improve the visibility of the symbols. (b) Peak amplitudes, *A*, of trajectories as defined in Fig. 2c for 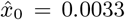. Amplitudes are decreasing for later peaks; the same trend has also been found for other values of 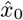 (data not shown). (c) Amplitudes of the first peak, *A*_1_, for all investigated initial positions, as long as the peak exists. (d,e) Normalised inter-peak distance, 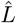, of trajectories as defined in Fig. 2c for all investigated 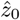 for (d) 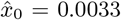 and (e) 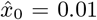. (f) Normalised inter-peak distance between the first and second peak, 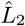, for 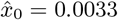 and 0.01. Lines are guides for the eyes.

The initial height of the particle, 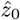, has a less pronounced effect on *N*. Trajectories with an initial position within the range 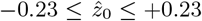 have the same number of peaks for identical values of 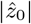 although the trajectories look different (*cf*. Fig. 2). For 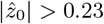, trajectories with 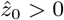 tend to have fewer peaks than those with 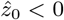 for the same magnitude of 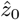. For the entire range of 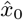, particles initially located at 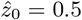 follow trajectories without any peaks.

#### 4.1.3 Peak amplitude of particle trajectories

The normalised peak amplitude, *Â* = *A*/(*W*/2), gives finer information about the shape of the trajectories. The data in Fig. 3b show the general trend that, for trajectories with more than a single peak, the amplitude of consecutive peaks is smaller than the amplitude of the previous peak, indicating that the spiralling motion of the particle is damped towards the outlet. The peak amplitudes are larger when particles are released at 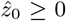, compared to trajectories with 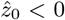 highlighting the influence of the vortex handedness on the particle’s motion in the junction under the investigated conditions.

Fig. 3c shows the behaviour of the normalised amplitude of the first peak, *Â*_1_, for all studied initial positions. For a given value of 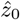, there is little change of *Â*_1_ with 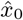, evidenced by the collapse of data for different values of 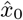, as long as the first peak exists. Changing 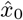 merely affects the number of peaks, but not the amplitude of the first peak. The amplitude *Â*_1_ increases with 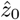 for negative and positive values of 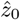, respectively, while the amplitude for 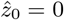 escapes the trend and defines a local maximum.

#### 4.1.4 Axial inter-peak distance

Like the amplitude of the peaks, *Â*, the axial distance between consecutive peaks, 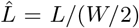, serves as a quantitative measure of the particle trajectories. We define 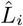 as the normalised axial distance (along the outlet axis) between the peaks *i* – 1 and *i*, as long as both peaks exist. The axial distance 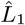 denotes the distance of the first peak from the inlet centre-line as illustrated in Fig. 2c.

Fig. 3d and 3e show results for 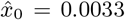 and 0.01, respectively. In both panels, only those curves are shown for which at least two peaks exist such that 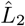 is defined. Since trajectories with 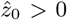 tend to have fewer peaks than those with 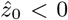, there are fewer data points for the former. In all observed trajectories, the inter-peak distance grows between consecutive pairs of peaks, 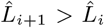.

Furthermore, 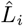 for a fixed *i* increases with 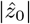: the farther away from the horizontal mid-plane a particle is located initially, the longer its spiralling trajectory is stretched toward the outlet. The data also show that 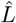 is larger for positive *z*_0_ than for negative *z*_0_ with the same magnitude 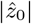. Comparing data from Fig. 3d and 3e, we see that 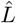 increases with 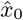: particles initially farther away from the vertical mid-plane have trajectories with larger inter-peak distances. Specifically, Fig. 3f shows the increase of 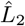 with 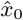.

#### 4.1.5 Residence time

The residence time of a particle in the junction (*cf*. Sec. 4.1.1) is an important parameter to consider. Due to the symmetry of the problem, a particle with 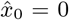 would in principle remain in the junction forever, *T*_res_ → ∞. For any finite value of 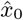, the residence time should be finite since the particle is located closer to one outlet and can leave the junction area more easily. In fact, Fig. 4a shows that *T*_res_ sharply decreases with 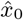 for small 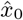 and drops further when 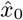 is increased.

**Figure 4:**
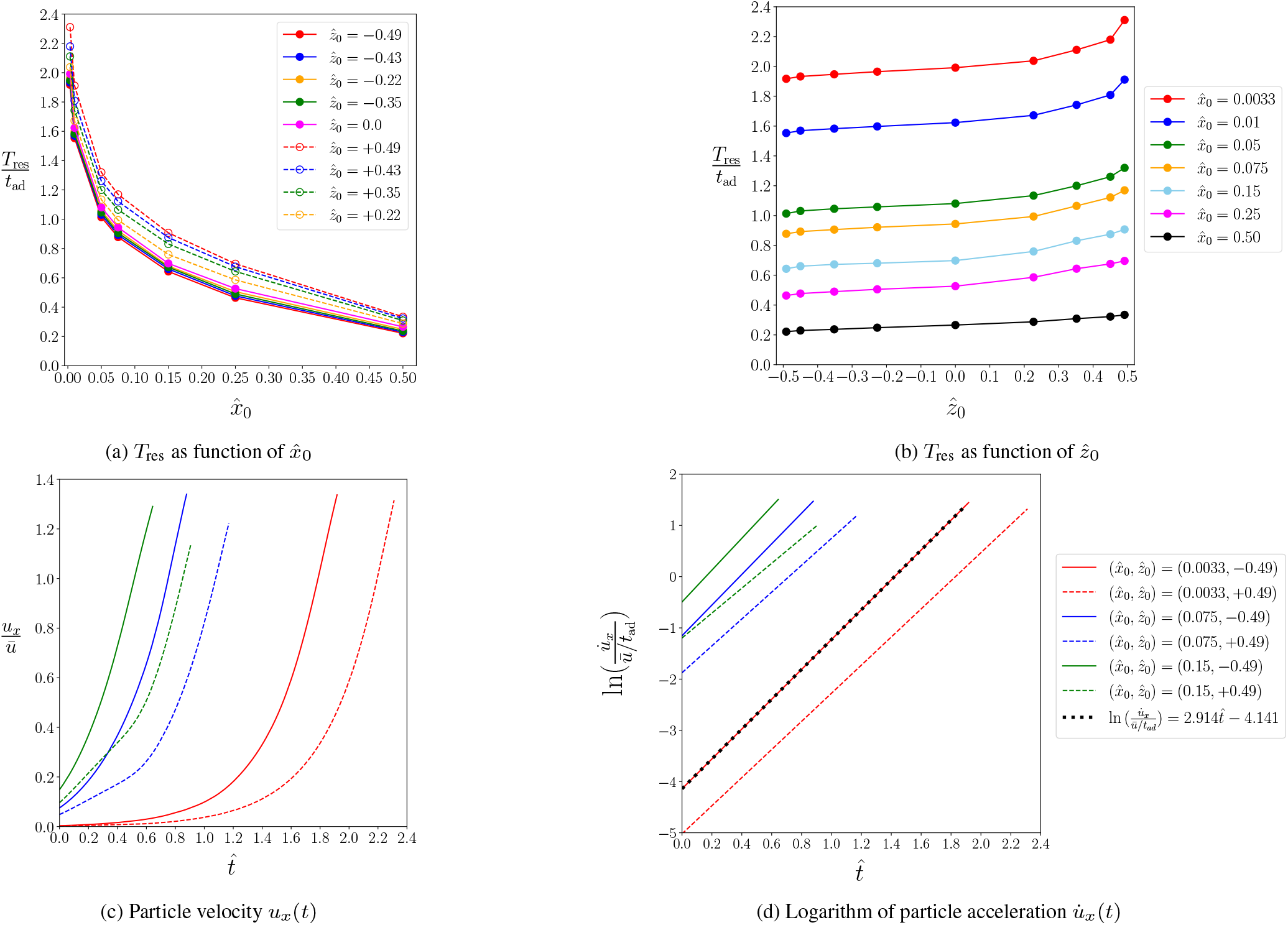
(a,b) Residence time, *T*_res_ (as defined in Sec. 4.1.1), as function of (a) 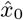 and (b) 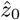. The residence time is normalised by the advection time, Eq. (5). Time evolution of (c) particle velocity and (d) logarithm of particle acceleration inside the junction along the outlet axis (*x*-axis).

Although *T*_res_ is largely determined by 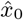, there is also a clear dependency on 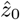 (Fig. 4b). For all studied values of 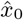, there is a mild increase of *T*_res_ with 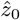: the residence time is shorter when the particle is initialised in the lower channel half 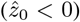 compared to initial positions in the upper channel half.

#### 4.1.6 Discussion

In the previous sections, we identified four clear trends upon increasing the initial position 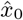: 1) the number of peaks, *N*, decreases, 2) the peak amplitude, *Â*_1_, is largely independent of 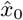, 3) the inter-peak distance, 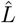, increases, and 4) the residence time, *T*_res_, decreases (*cf*. Fig. 3a, Fig. 3c, Fig. 3d and 3e, and Fig. 4a, respectively). When changing the initial position 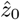 instead, we observed that 1) the number of peaks is smaller for 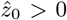 than for 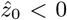 for the same magnitude of 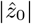, 2) the peak amplitude increases with 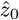, 3) the inter-peak distance is smaller for 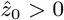 than for 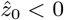 for the same magnitude of 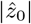, and 4) the residence time mildly increases with 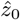 (*cf*. Fig. 3a, Fig. 3c, Fig. 3d and Fig. 4b, respectively).

In order to understand the behaviour of the residence time, we first analyse the velocity and acceleration of the particle along the outlet axis (*x*-axis) inside the junction (Fig. 4c and 4d). Particles that are initially close to the vertical mid-plane (small 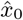) accelerate more slowly than a particle that enters the junction farther away (larger 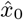). The slow acceleration of a particle with small 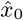 is caused by the particle covering nearly identical portions of volume on either side of the mid-plane; hence the net force accelerating the particle towards one of the outlets is nearly zero. Once the particle is slowly moving towards one outlet, it gets increasingly exposed to the drag force exerted by the flow in that outlet channel, and the particle acceleration, and therefore velocity, increases. A particle that enters the junction far away from the mid-plane, on the one hand, is closer to the outlet and, on the other hand, experiences a larger drag force early on and leaves the junction sooner; hence, the residence time is smaller. The residence time also depends on 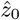 (Fig. 4b), but to a lesser extent than it depends on 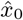. This dependency stems from the particle acceleration 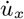 being smaller for particles entering the junction above the horizontal mid-plane 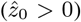. The mechanism behind this effect will become obvious at the end of this section. Interestingly, the acceleration increases exponentially with time as revealed in Fig. 4d.

The behaviour of the number of peaks, *N*, and the inter-peak distance, 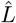, is determined by an interplay of the residence time and the ability of the particle to follow the vortex and revolve about the outlet axis (*x*-axis). The slower the particle moves towards the outlet and the faster it revolves about the outlet axis, the larger the number of peaks and the smaller the inter-peak distance. Fig. 5a depicts the distribution of the vorticity magnitude of the unperturbed flow field in the junction on the horizontal mid-plane 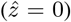. The vortex can be recognised by a region of high vorticity stretched along the outlet axis. The vortex causes the particle to revolve about the outlet axis with an angular velocity Ω_*x*_, thus creating the spiralling trajectories illustrated in Fig. 2a. Fig. 5b shows the angular velocity, averaged over the time the particle is located inside the junction area, 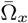. The number of peaks *N*, residence time *T*_res_ and the average angular velocity 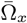 are related according to

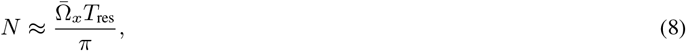

remembering that a full revolution of 2*π* radians leads to two peaks as defined in Fig. 2c. Particles initially located closer to the vertical mid-plane (small 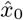 experience a higher angular velocity than particles entering the junction farther away from the mid-plane (large 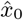). Combining the faster revolution 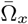 with the smaller linear acceleration 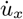 (and therefore larger residence time) for particles with 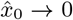 explains why these particles have the largest number of peaks, *N*, according to Eq. (8). Since peaks are only counted in the junction area which has a fixed size, a trajectory with more peaks has smaller inter-peak distances 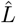, which explains the trend of 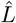 becoming smaller for 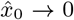. Looking at particles with small 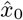, Fig. 5b reveals that 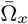 is significantly smaller for particles with 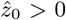, while the residence time in Fig. 4b shows only a moderate increase for these particles, compared to particles with 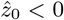. Consequently, particles with 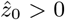 show fewer peaks than particles with 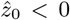 as long as 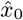 is small. The smaller revolution frequency of particles with 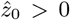 can be understood as follows: Fig. 5c shows that, upon entering the junction, these particles move over the vortex, rather than getting trapped in the vortex, then reach a region with relatively small vorticity, before finally entering the vortex. Averaged over the time they spend in the junction, these particles, therefore, do not revolve as much as the particles that are able to enter the vortex immediately.

**Figure 5:**
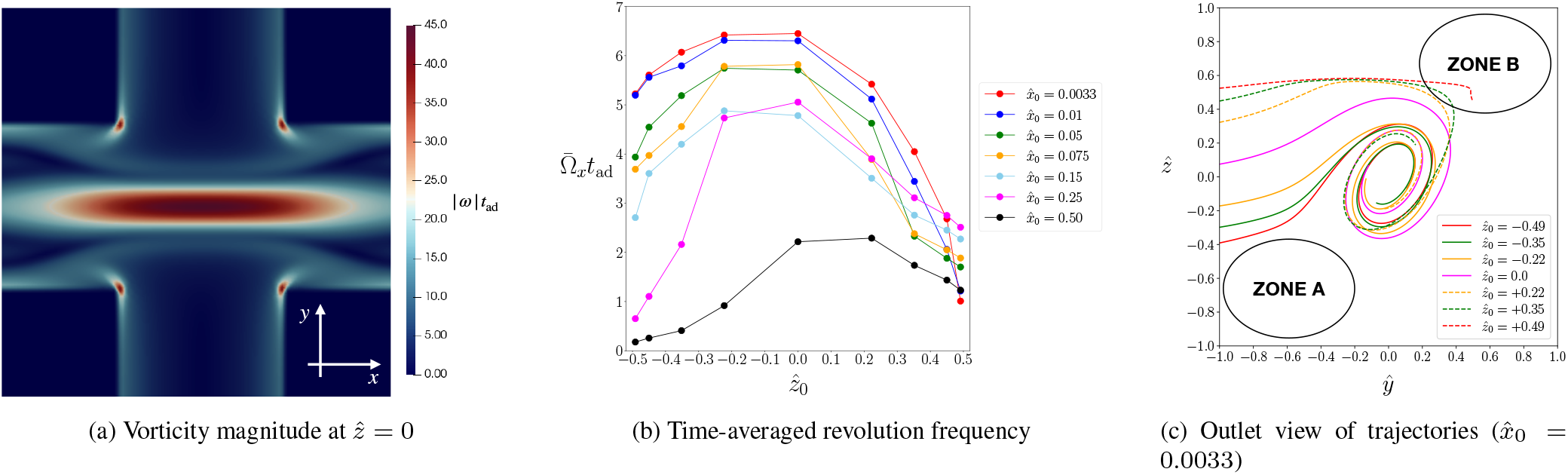
(a) Dimensionless vorticity magnitude of the unperturbed flow, *ωt*_ad_ on the horizontal mid-plane, 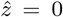. (b) Dimensionless time-averaged angular velocity, 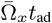, of the particle revolving about the outlet axis for different initial positions. (c) Outlet view of particle trajectories for 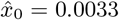. Zones A and B indicate regions with small flow velocity magnitude (*cf*. Fig. 1d).

In order to understand the variation of the peak amplitude with the initial positions 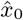 and 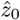, we first inspect the particle trajectories as seen from the outlet (projection on the inlet-height plane, Fig. 5c). A particle that is released at 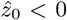 experiences an immediate upward lift force and is drawn into the vortex where the particle revolves with small amplitude. Contrarily, a particle with larger 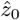 features a flatter trajectory, tends to avoid the vortex, and approaches low-velocity zone B. Only after the encounter with zone B, the particle is pulled into the vortex, resulting in a larger peak amplitude *Â*_1_. None of the particles simulated here are able to reach zone A; this would only be possible for particles with very small confinement *χ* released at 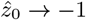. The different shapes of particle trajectories can be understood by inspecting the streamlines in Fig. 1d: streamlines with slightly negative 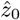 turn upwards and reach the vortex immediately, while streamlines with slightly positive 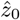 tend to avoid the vortex and approach zone B; only streamlines near the bottom of the channel are drawn towards zone A. The independence of *Â*_1_ from 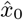 can be explained by the shape of the high-vorticity region in Fig. 5a. Since the vorticity profile is largely constant along the outlet axis inside the entire junction area, the initial interaction between the particle and the vortex is not strongly sensitive to 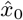.

Finally, Fig. 5c explains the trend of the residence time to increase with 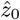 (Fig. 4b). Particles with larger 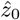 spend some time in the low-velocity zone B, which delays the particle’s acceleration towards the outlet, therefore increasing the residence time in the junction.

### 4.2 Effect of particle confinement

In this section, we analyse the effect of the size of the particle on its dynamics in the cross-slot junction. We investigate five different confinement values, *χ* ∈ [0.35, 0.65] and the limiting case of a mass-less point particle, *χ* → 0. In order to keep the number of free parameters manageable, we focus on 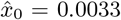, which leads to the largest number of peaks, and 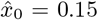, which is the lowest value of 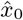 for which nearly no peaks have been observed in Fig. 3a. Additionally, we fix the initial positions at 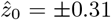 which is the lateral equilibrium position of a particle with *χ* = 0.65 in a straight channel with the same dimensions and Reynolds number as the inlet channel of the cross-slot geometry. The hydrodynamic conditions and channel aspect ratio are kept constant at *Re* = 120 and *α* = 0.6, respectively.

#### 4.2.1 Initial position at 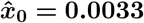

In Sec. 4.1, we found that the vortex has a more profound influence on the particle dynamics when 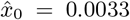 than for the other investigated initial positions. Fig. 6 shows the particle trajectories in the junction for the investigated confinement values for 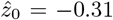 in (a,e) and 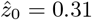 in (b,f). As the confinement %increases, the characteristic spiral trajectories seen for particles with low confinement are less pronounced or no longer observed at all. For 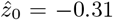, the particle with *χ* = 0.65 leaves the junction without any oscillation. For 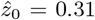, no particle with *χ* ≥ 0.5 oscillates in the junction. A mass-less point particle, which follows the fluid streamlines, shows the most oscillatory behaviour of all particles for 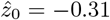; contrarily, when initialised at 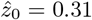, the point particle avoids the vortex and hardly oscillates.

**Figure 6:**
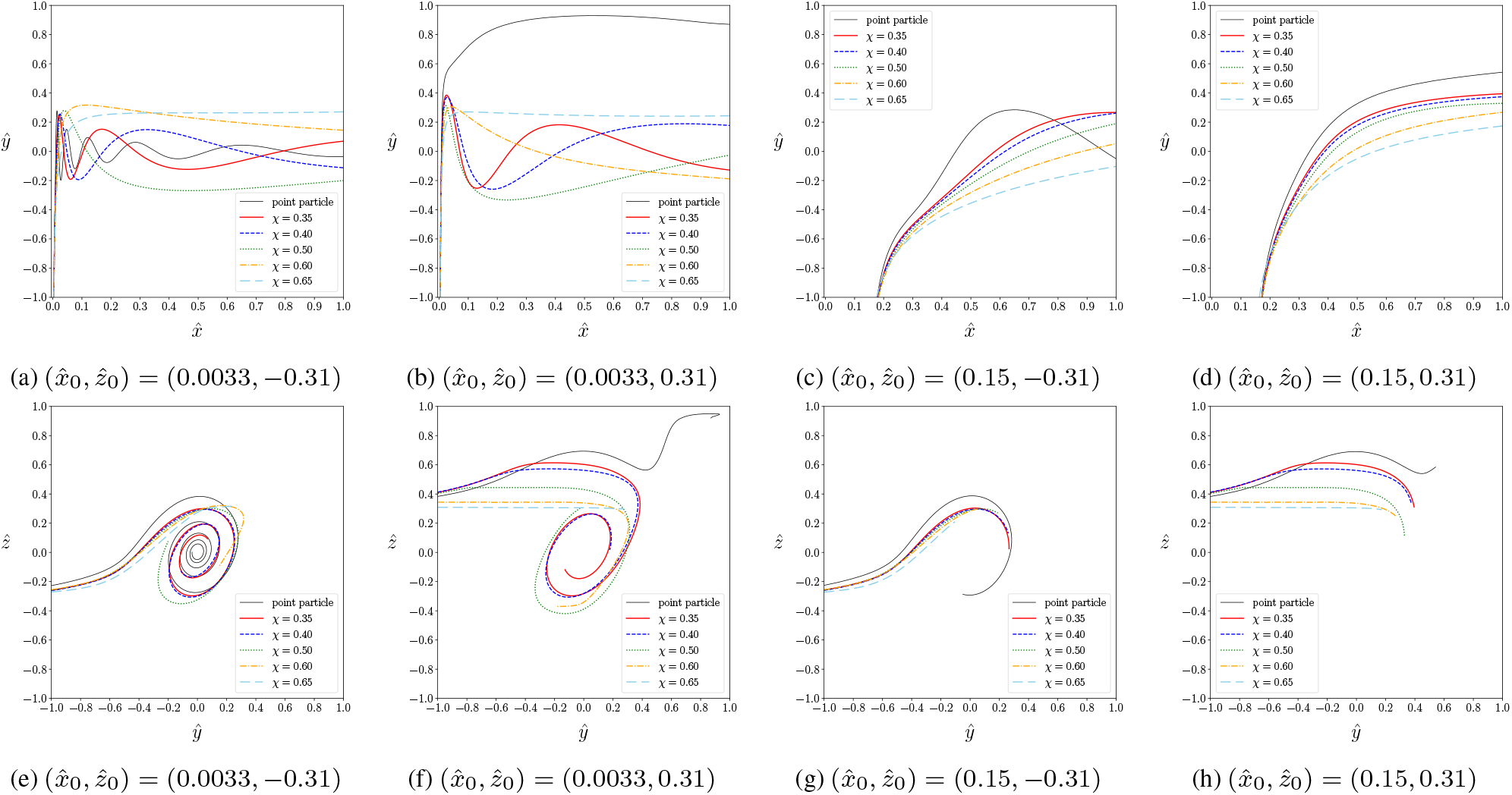
Particle trajectories for different confinement values *υ*;. (a–d) Top views and (e–h) outlet views. The initial particle position is (a,e) 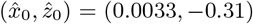, (b,f) 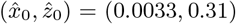 (c,g) 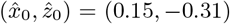 and (d,h) 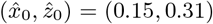. The trajectories in (e–h) are shown up to the point where the particle reaches the outlet channel at 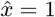.

The revolution frequency of the particle about the outlet axis, Ω_*x*_, provides a complementary view. When particles enter the junction below the horizontal mid-plane, 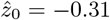 (Fig. 7a), Ω_*x*_ is highest for particles with *χ* < 0.5, including the point particle, and Ω_*x*_ decays quickly for larger confinement. For 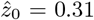 (Fig. 7b), particles with 0.35 ≤ *χ* < 0.5 show a behaviour similar to that for 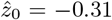. In contrast, both the point particle and the particles with *χ* ≥ 0.5 hardly oscillate when released above the horizontal mid-plane, 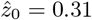. While the oscillation of the point particle is suppressed by the particle reaching the low-velocity zone B, larger particles are strongly confined when they approach the top wall of the junction and, hence, are unable to revolve freely.

**Figure 7:**
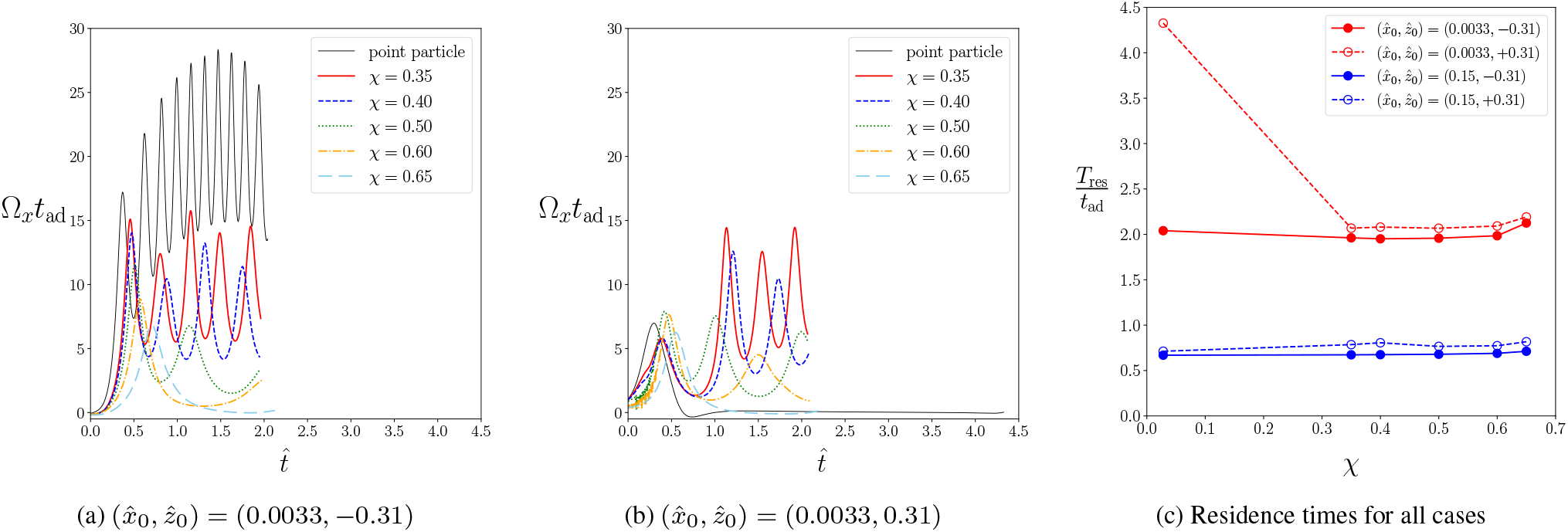
Non-dimensional revolution frequency Ω_*x*_ for different confinement values *χ* for initial positions (a) 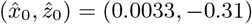 and (b) 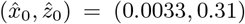. (c) Non-dimensional residence time *T*_res_ for different confinement values *x* and all studied initial positions. Each curve in (a,b) is shown for its range [0, *T*_res_] during which the particle is located in the central junction area, thus each curve has a different range.

Fig. 7c displays the residence time, *T*_res_, for all investigated cases. For particles that are released below the horizontal mid-plane, 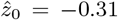, confinement has a negligible effect on the residence time: for all particles, including the point particle, *T*_res_ ≈ 2*t*_ad_. The insensitivity of the residence time to the confinement can be explained by the observation that all particles remain in the vortical zone throughout their passage (*cf*. Fig. 6e) where particles are efficiently advected towards the outlet. The situation is different for 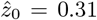 where the residence time strongly depends on particle size. While particles with 0.35 ≤ *χ* ≤ 0.5 leave the junction nearly as quickly as their respective counterparts at 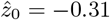, both the point particle and the larger particles (*χ* > 0.5) require more time to leave the junction area. The increased residence time of the point particle is caused by the point particle reaching and remaining in the low-velocity zone B as seen in Fig. 6f. The large particles with *χ* > 0.5 do not reach zone B; instead, they are nearly in contact with the top wall, which reduces their ability to follow the flow around the vortex or towards the outlet.

#### 4.2.2 Initial position at 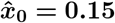

In Sec. 4.1, we demonstrated that the vortex has little effect on the motion of the particle when the particle enters the junction farther away from the vertical mid-plane, for example at 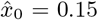. Here, we show that, even if particle confinement is varied, the vortex continues to have little influence on the motion of the particle at 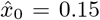. Fig. 6 depicts the particle trajectories in the junction for the investigated confinement values for 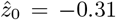 in (c,g) and 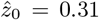 in (d,h). For both 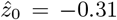 and 0.31, the particle experiences a strong acceleration towards the outlet once it enters the junction area since most of the particle volume is located in the outlet-facing half of the junction. During its passage, the particle does not reach the vortex core and, therefore, does not undergo a strong revolution about the outlet axis (Ω_*x*_ not shown for 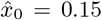). Consequently, in nearly all situations, the particle leaves the junction without displaying a peak in its trajectory. The only exception is the point particle for 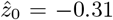 with a single peak, although the point particle essentially follows a similar trajectory as the finite-size particles.

The residence time *T*_res_ is consistently lower for 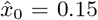 compared to 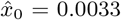 (Fig. 7c). The shorter residence time is caused by the particle with 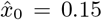 being accelerated towards the outlet earlier and, at least to some extent, the shorter distance to travel to the outlet. The residence time is essentially independent of confinement *χ*, and it is nearly the same for 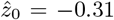 and 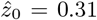: all particles follow roughly the same trajectory and therefore experience a similar acceleration along the outlet axis (Fig. 6).

## 5 Conclusions

We used the immersed-boundary-lattice-Boltzmann method to numerically investigate the dynamics of a rigid neutrally-buoyant spherical particle in a cross-slot junction for a channel height-to-width ratio of 0.6 and a Reynolds number of 120 for which a steady vortex exists in the junction. We characterised the effects of the initial position and the particle confinement *χ* = 2*α*/*H* (where *a* is the particle radius and *H* is the height of the channel) on the particle trajectories in the junction. Particles generally show a swirling motion within the cross-slot. Thus, observed trajectories were described in terms of macroscopic observables, such as the number of peaks of each trajectory (*N*), the peak amplitudes (*A*), the axial inter-peak distances (*L*), and the residence time within the junction area (*T*_res_). The simulations also provide access to microscopic observables, such as instantaneous linear and angular velocity and acceleration, which were used to rationalise the particle’s behaviour.

There are several key observations when varying the location at which the particle with fixed size enters the junction:

- The residence time *T*_res_ strongly depends on the initial position of the particle. A particle entering the junction closer to the vertical mid-plane is accelerated more slowly towards its closest outlet, thus leading to a larger residence time. There is a mild effect of the initial height of the particle on the residence time. The behaviour of the residence time can be explained by the interaction of the particle with low-velocity areas in the flow field.
- Two competing mechanisms determine the number of peaks *N* and the axial inter-peak distance *L* seen in the particle trajectories: particles that are able to follow the vortex and revolve about the outlet axis more easily show a larger number of peaks and shorter inter-peak distances. Contrarily, particles with a shorter residence time have a trajectory with fewer peaks and larger inter-peak distances. As a net effect, the number of peaks and inter-peak distance depend strongly on the lateral position and weakly on the height position of the particle when entering the junction.
- The amplitude *A* of consecutive peaks of a given trajectory generally decrease as the particle follows the dissipating vortex along the outlet channels. The initial height of the particles has a strong effect on the amplitude: depending on the initial position, the particle either moves along the vortex, resulting in a larger amplitude, or it moves against the vortex, resulting in fast migration towards the centre of the junction and a smaller amplitude.

The properties of the particle trajectories also depend on the particle confinement:

- When entering the junction close to the vertical mid-plane, larger particles have a slightly higher residence time *T*_res_ since a large fraction of the particle is initially located in the region drawn into the second outlet.
- The trajectories of particles with small confinement show more peaks *N* with smaller inter-peak distance *L* because smaller particles are able to revolve faster about the outlet axis.
- Particles with larger confinement show a higher amplitude *A* of the first peak since these particles tend to pass the vortex first before being drawn into the vortex.

The cross-slot geometry has been successfully employed in flow cytometry to deform cells [30, 31, 51] where cell deformation is used as a bio-marker for the cell state [39]. In deformability cytometry using a junction, cells or particles are usually inertially focused before entering the junction [31]. Therefore, cells or particles are expected to enter and traverse the junction with typical trajectories which can be quantified using the metrics used in the present study. The videos of Gossett et *al.* [31] display polystyrene beads, droplets and cells performing an oscillating trajectory in the cross-slot junction as seen from above (along the *z*-axis), similar to the ones we observed in Sec. 4.1. Zhang et al. [50] reported oscillations in the trajectory of glass beads in a cross-slot channel with square cross-section at *Re* = 80. The trajectories of the rigid particle in our study show a good qualitative agreement with those trajectories seen under steady-vortex conditions published previously [31, 50].

Our study provides a deeper understanding of the dynamics of a rigid particle interacting with the vortex in a cross-slot junction at moderate inertia. The macroscopic observables (*N, A, L, T*_res_) used in our work might help distinguish between particles with different properties in experimental cross-slot flows. To this end, a future investigation of the effect of channel geometry, in particular aspect ratio, and Reynolds number appears beneficial. Since most inertial microfluidic applications involve deformable cells, a study of deformable particles in cross-slot flows would provide further insight into the particle dynamics of real-world applications.

## Acknowledgements

T.K. received funding from the European Research Council (ERC) under the European Union’s Horizon 2020 research and innovation program (803553). This work used the Cirrus UK National Tier-2 HPC Service at EPCC (https://www.cirrus.ac.uk). The authors thank Emmanouil Falagkaris for contributions to the code development.

For the purpose of open access, the authors have applied a Creative Commons Attribution (CC BY) licence to any Author Accepted Manuscript version arising from this submission.

## Conflict of interests

H.T. is co-founder and CTO of Cytovale Inc., a company which is commercialising a deformability cytometry technology based on cross-slot flows. D.D. is co-founder and on the Board of Directors of Cytovale Inc. L.G. was employed by Cytovale Inc. during the preparation of the manuscript. T.K. has received financial support from Cytovale to partially fund a PhD student who is not a co-author of this paper. The remaining authors have no conflict of interest to declare.

## Author contributions

T.K. acquired funding, conceptualised and oversaw the study. B.O. validated the code. K.K. generated the data, carried out the analysis and wrote the original manuscript. All authors discussed the results and edited the manuscript.

## Data availability

The data that support the findings of this study are available from the corrensponding author upon reasonable request.

## A Lateral equilibrium positions in straight channels

### A.1 Test case: effect of initial position and confinement on lateral migration in square duct

We determine the lateral equilibrium position of a rigid neutrally buoyant spherical particle in a Poiseuille flow through a straight duct with square cross-section at *Re* = 100. The duct has a width and height *W* = 35Δ*x*. The particle radius is varied such that confinement values *χ* = 0.1, 0.2 and 0.2857 are obtained. Periodic boundary conditions are used at the inlet and outlet of the duct. With *L* = 6 W, the domain length is sufficiently long to ensure that periodic images of the particle do not interact. The viscosity is set to *ν* = 1 /30Δ*x*^2^/Δ*t*.

We compare the equilibrium positions found using our IB-LB solver with those reported by Lashgari *et al*. [48] who originally proposed this case. Fig. 8 shows excellent agreement of the equilibrium positions. As expected, equilibrium positions are independent of the initial position. Larger particles assume equilibrium positions closer to the duct centre at the mid-point of a duct edge.

**Figure 8:**
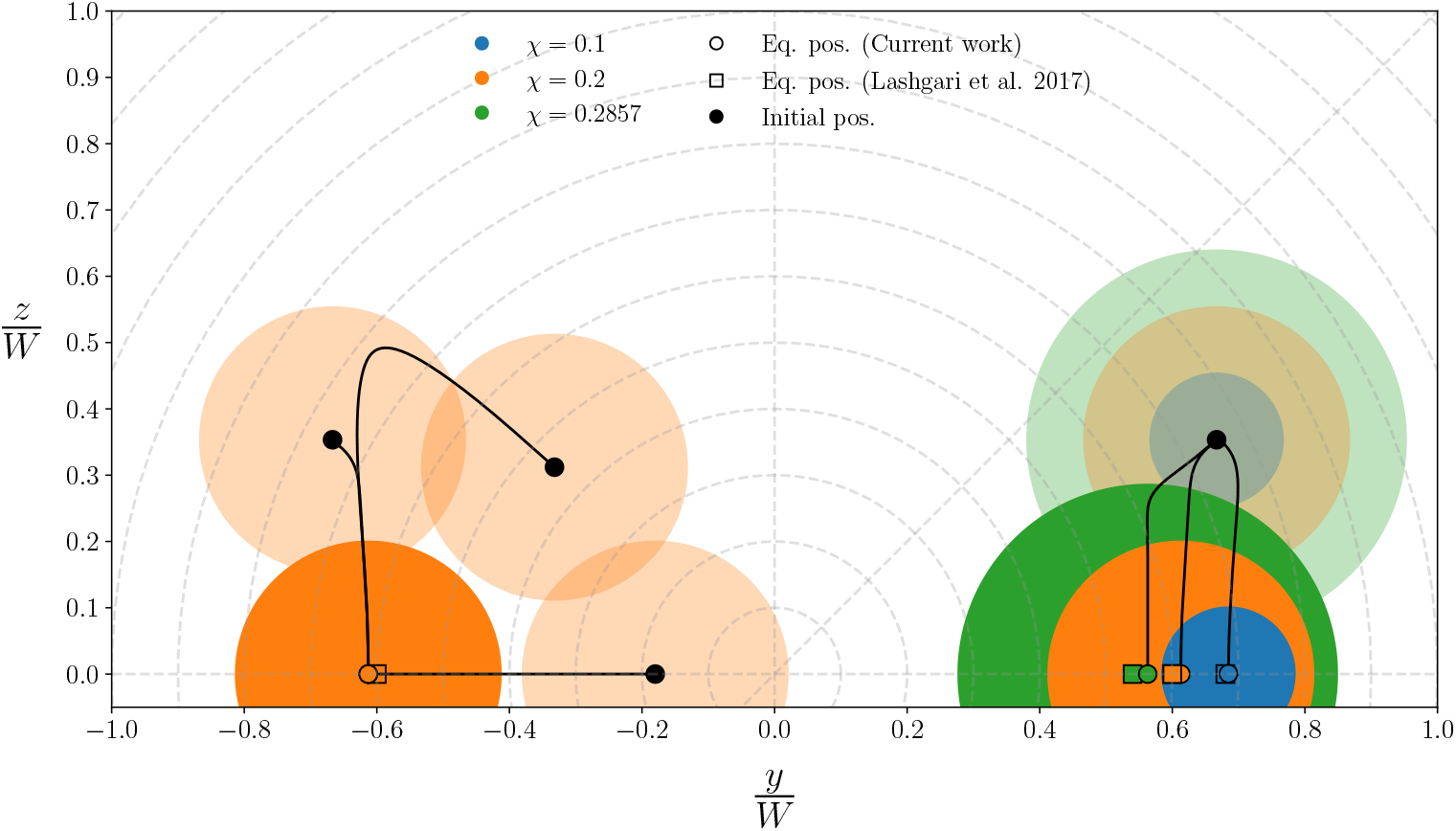
Results of the test case in App. A.1. Particles move along the *x*-axis and migrate in the *y-z*-plane which is the cross-section of the square duct. The origin of the coordinate system marks the centreline of the duct. Left: migration paths and lateral equilibrium positions of particles with different initial position at *χ* = 0.2. Right: migration paths and lateral equilibrium positions of particles with the same initial position and different *χ*. Lateral equilibrium positions are compared to those reported by Lashgari *et al.* [48].

### A.2 Equilibrium positions in a straight channel equivalent to the inlet of the cross-slot

We perform simulations of a rigid neutrally buoyant spherical particle flowing through a straight channel (along the *x*-axis) with the same cross-sectional shape as the inlet of the cross-slot geometry to identify the lateral equilibrium positions for a given particle size. The simulation parameters are the same as in Sec. 3, in particular, the Reynolds number is *Re* = 120 and the channel aspect ratio is *α* = 0.6. Periodic boundary conditions are used at the inlet and outlet of the duct. With *L* = 4*W*, the domain length is sufficiently long to ensure that periodic images of the particle do not interact [52]. The schematic in Fig. 9a illustrates the initial configuration of the particle in the channel.

**Figure 9:**
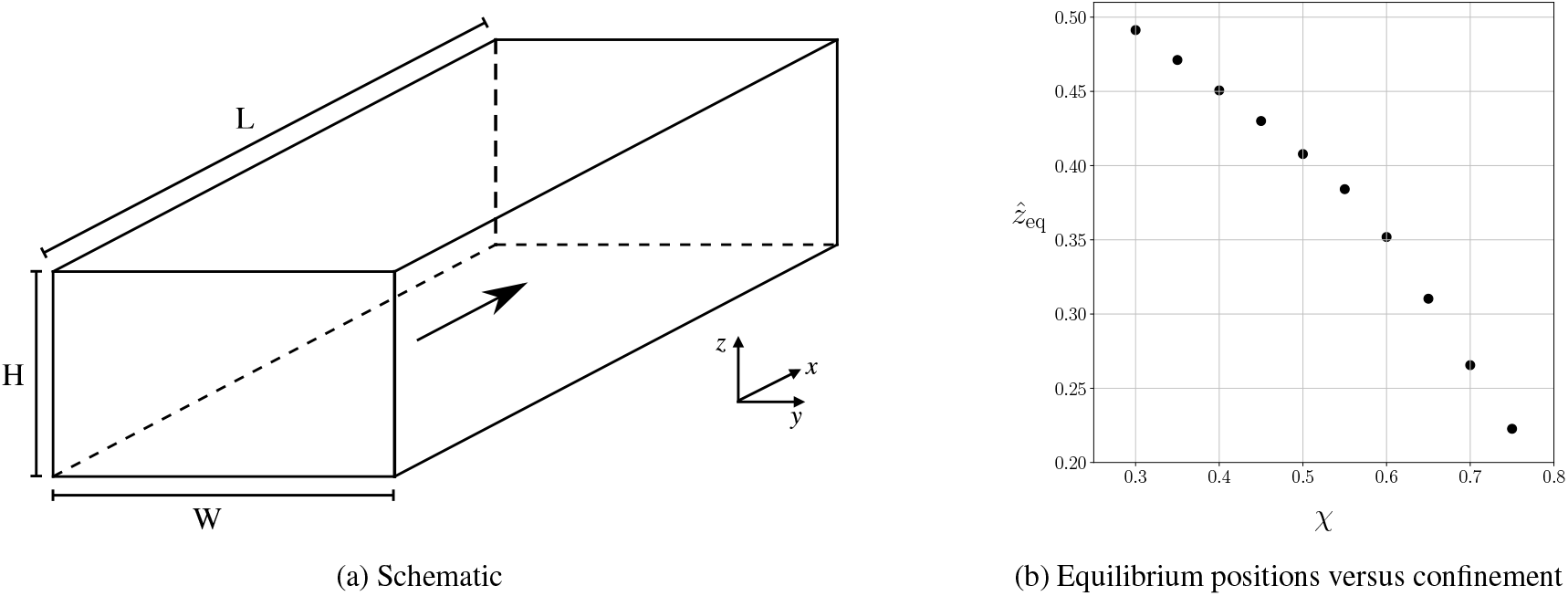
Simulation of a rigid neutrally-bouyant spherical particle in a Poiseuille flow through a straight rectangular channel with the same aspect ratio as the inlet of the cross-slot geometry. The particle confinement *χ* is varied in the range [0.3, 0.75]. (a) Schematic of the set-up. The particle is initialised on the grey plane which is located midway between the side walls. (b) Equilibrium positions 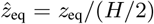 as function of confinement *χ*. The particle remains on the mid-plane (*y* = 0) at all times.

Fig. 9b shows the identified equilibrium positions *z_eq_* that we used as the initial positions for our main study in Sec. 4. Our simulations confirm the known observation that, for the studied parameter values, the equilibrium position of the particle is located mid-way along the long cross-sectional axis (*y* = 0) [53]; the particle remains on this mid-plane during its lateral migration.

